# Inferring the Spiking Rate of a Population of Neurons from Wide-Field Calcium Imaging

**DOI:** 10.1101/2020.02.01.930040

**Authors:** Merav Stern, Eric Shea-Brown, Daniela Witten

## Abstract

Wide-field calcium imaging techniques allow recordings of high-resolution neuronal activity across one or more brain regions. However, since the recordings capture light emission generated by the fluorescence of the calcium indicator, the neural activity that drives the calcium changes is masked by the calcium indicator dynamics. Here we develop and evaluate new methods to deconvolve the calcium traces and estimate the underlying neural spiking rate. Our methods take into account both the noise in the recordings and the temporal dynamics of the calcium indicator response. Our first proposal estimates firing rates that are constant over discrete time bins. The size of each time bin depends on the data and is determined dynamically. Our second proposal estimates the rate as a continuous function and is meant for studies that look for slow rate fluctuations rather than abrupt changes. We compare our results with those of two alternative approaches: direct deconvolution using a ‘first differences’ approach, and the ‘Lucy-Richardson’ image recovery method, adapted to recover temporal dynamics. We show that our methods outperform competitors on synthetic data as well as on wide-field calcium recordings in which the spikes were recorded in parallel using multi-channel silicon probe.

## 1 Introduction

Recent developments in optogenetics allow for the recording of high-resolution images of neuronal activity from the entire dorsal surface of the cortex in behaving animals (Makino et al. 2017, Chen et al. 2017, Allen et al. 2017), through the use of fluorescent calcium indicator molecules (Chen et al. 2013). Each pixel captures the fluorescence arising from dozens to thousands of neurons. Images can be steadily collected for hours at a high rate, from dozens to hundreds of Hertz. When the images are aggregated over time, a fluorescence trace can be extracted for each pixel. Inferring the time-varying neuronal activity from a given fluorescence trace is a challenging deconvolution problem.

Wide-field recording techniques date back a few decades. Before optogenetic manipulations were available, wide-field techniques used a combination of a single photon microscope and a camera to record the natural changes in illumination due to hemodynamic changes (Masino et al. 1993, for example). Because wide-field recordings cover large fields of view, the luminescence captured in each pixel originates from multiple sources (blood vessels and/or neurons). Hence the luminescence is strong enough to be captured through the skull, when the skull is relatively thin, as is the case for mice. This avoids invasive procedures, which in turn allows recordings of *in vivo* brain activity in behaving animals. It is also possible to conduct wide-field recordings in anesthetized animals (Kalchenko et al. 2014) or *in vitro* slices.

In recent years, the development of optogenetics has enabled neuronal recordings in *vivo* for long periods of time and across multiple days by combining optogenetic manipulations, which were originally developed for two photon microscopy, with chronically implanted windows that expose large fields of view. Furthermore, wide-field imaging can now be used to image neural activity across multiple brain regions (Silasi et al. 2016). As a result, wide-field imaging has rapidly become a standard recording technique (Clancy et al. 2019, Mann et al. 2017, Musall et al. 2018, Chen et al. 2017, Allen et al. 2017, Aimon et al. 2015, Wekselblatt et al. 2016, Makino et al. 2017). However, suitable approaches for deconvolution of wide-field calcium recordings are notably absent. In this paper, we develop a statistical model to extract mesoscale neural activity from wide-field recordings.

The problem of inferring the underlying neuronal activity from a fluorescence trace has recently been considered by a number of authors, in the case of fluorescence traces that result from the activity of a single neuron (Jewell & Witten 2018, Jewell et al. 2019, Friedrich et al. 2017, Pnevmatikakis et al. 2016). These papers make use of an auto-regressive model, originally proposed in Vogelstein et al. (2009), that associates the fluorescence *y_t_* of a single neuron at the *t*th timepoint with the unobserved calcium *c_t_* at the *t*th timepoint,

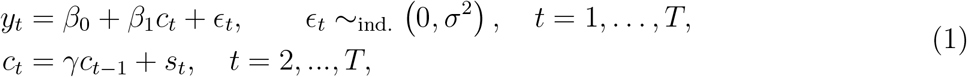

where *s_t_* ≥ 0 allows for the occurrence of a spike at the *t*th timepoint, and where *γ* ∈ (0,1) is the rate of calcium decay. In words, (1) indicates that the calcium decays exponentially over time, unless there is a spike at the tth timepoint, in which case it increases; furthermore, the observed fluorescence is a noisy realization of the underlying calcium at each timepoint. In (1), *β*_0_ corresponds to the baseline fluorescence, which must be estimated; however, we can set *β*_1_ = 1 without loss of generality, as this simply amounts to scaling the calcium by a constant factor. To fit the model (1) with *β*_1_ = 1, Friedrich et al. (2017), Jewell & Witten (2018), and Jewell et al. (2019) solve the optimization problem

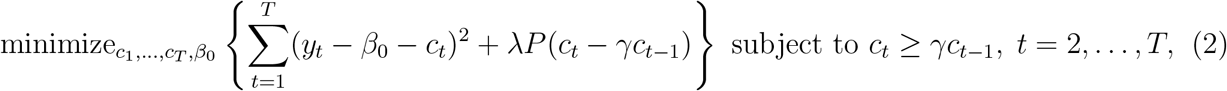

where λ is a nonnegative tuning parameter, and where *P*(·) is a penalty function designed to induce sparsity in its argument — that is, to encourage *c_t_* – *γc*_*t*−1_ = *s_t_* = 0 — so that at most timepoints, no spike is estimated to occur. Friedrich et al. (2017) makes use of an *ℓ*_1_ penalty (Tibshirani 1996), while Jewell & Witten (2018) and Jewell et al. (2019) instead use an *ℓ*_0_ penalty.

However, fluorescence traces that result from wide-field calcium imaging recordings correspond to the activity of a collection of neurons rather than the activity of a single neuron. Consequently, single-neuron deconvolution solutions, which assume that at most time points there are no spikes, cannot be directly applied. In this manuscript, we propose to extend the model (1), and the corresponding optimization problem (2), to the setting of wide-field calcium imaging recordings. We apply this new approach for deconvolution of wide-field calcium imaging recordings to data from the retrosplenial cortex (Swanson et al. 2018).

## 2 A Model for Wide-Field Recordings and Neuronal Activity

### 2.1 Extension of the Model (1) to Wide-Field Recordings

To begin, we consider the model (1), in a setting in which the observed fluorescence trace is the sum of the fluorescences associated with each of *p* neurons recorded at a given pixel. For the *j*th neuron, *j* = 1,…, *p*, (1) takes the form

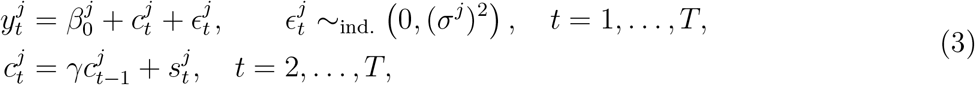

where we assume that the rate of calcium decay, *γ*, is the same for all *p* neurons. However, in the wide-field case we do not separately observe the fluorescence for each of the *p* neurons; we instead observe their summed fluorescence. Summing (3) across the *p* neurons yields the model

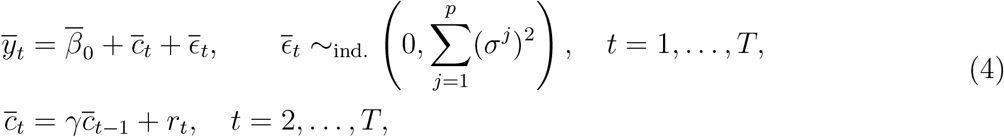

where 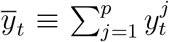 is the total observed fluorescence of the *p* neurons, and where 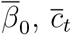, and 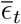 are the total baseline fluorescence, total calcium at the *t*th timepoint, and total noise at the *t*th timepoint, respectively. In (4), we can interpret *r_t_* as the amount that the calcium increases, at the *t*th timepoint, as a result of spiking events in the *p* neurons; we will refer to this as the *spiking rate.* Because *p* is potentially quite large, on the order of hundreds to thousands of neurons, we do not expect *r_t_* to be sparse. However, critically, we *do* expect *r_t_* to (for the most part) take on similar values at nearby timepoints: we do not expect the rate to vary over time in an arbitrary way. We will explore this point in greater detail in the next section.

The total baseline fluorescence, 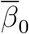, typically depends on the processing performed on the observed fluorescence; see e.g. the Δ*F/F* pre-processing of Chen et al. 2017.

### 2.2 Optimization Problem

The model (4) leads naturally to the optimization problem

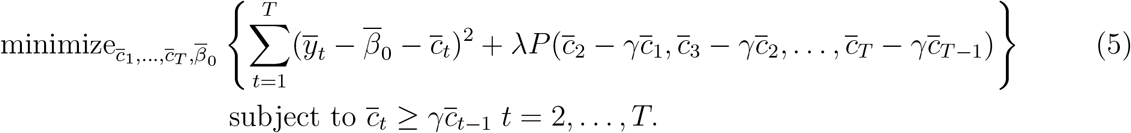

Problem (5) closely resembles the optimization problem (2) used to deconvolve the fluorescence trace for a single neuron (Friedrich et al. 2017, Jewell & Witten 2018, Jewell et al. 2019). For that task, the authors considered the use of a sparsity-inducing penalty, because a neuron is not expected to spike at most timepoints. By contrast, here we are considering a model in which 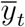 represents the total fluorescence at the *t*th timepoint summed over *p* neurons, and in which 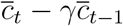 represents the spiking rate at the *t*th timepoint across all *p* neurons. Therefore, in the context of wide-field imaging, we do not want *P*(·) to be a sparsity-inducing penalty. Instead, we want *P*(·) to encourage adjacent timepoints to, for the most part, have similar values of *r_t_*.

For convenience, in what follows, we will reparametrize (5) in terms of *r*_1_,…, *r_T_*, where 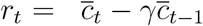 for *t* = 2,…, *T* as defined in (4), and where 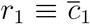. The latter is strictly a definition intended for notational convenience. This reparametrizaton can be also expressed in matrix form, 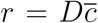, where *D* is a *T* × *T* full-rank matrix with 1’s on the diagonal and –*γ*’s just below the diagonal. This leads to a rephrasing of the optimization problem (5) as follows:

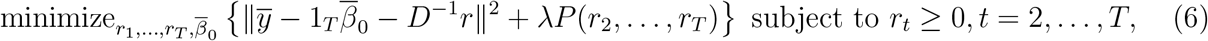

where 1_*T*_ is a vector of length *T* with all elements equal to 1.

#### Proposition 1.

*The optimization problem* (5) *is equivalent to* (6), in *the sense that* 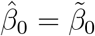 *and* 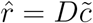, *where* 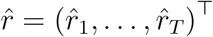 and 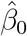 *solve* (6), *and* 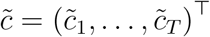 *and* 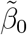 *solve* (5).

We propose two possible forms for *P*(·):

1. *Dynamically-Binned Spiking Rate.* We consider the penalty

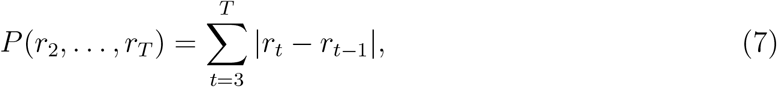

which is a *fused lasso,* or *total variation denoising,* penalty (Tibshirani et al. 2005, Tibshirani et al. 2012, Condat 2013). This penalty encourages the spiking rate, *r_t_*, to be constant over time, with only occasional changepoints, as shown in Figure 1B. This will yield estimates of the spiking rate that are constant within a bin, where the bins are themselves adaptively estimated from the data.
2. *Continuously-Varying Spiking Rate*. We consider the penalty

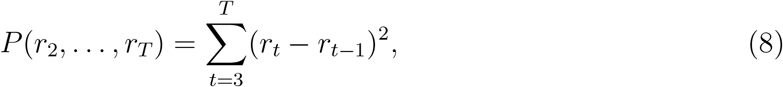

which encourages *r_t_* ≈ *r*_*t*−1_, so that the spiking rate varies continuously over time, as shown in Figure 1C. For simplicity, we can represent both penalties as

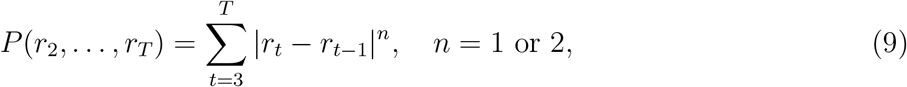

where *n* =1 corresponds to dynamically-binned (7) and *n* =2 corresponds to a continuously-varying rate (8). The optimization problem (6) with penalty (9) can be written as

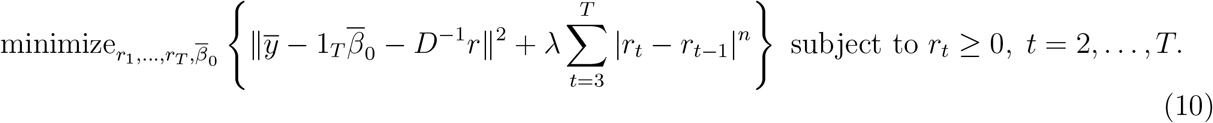

**Figure 1:**
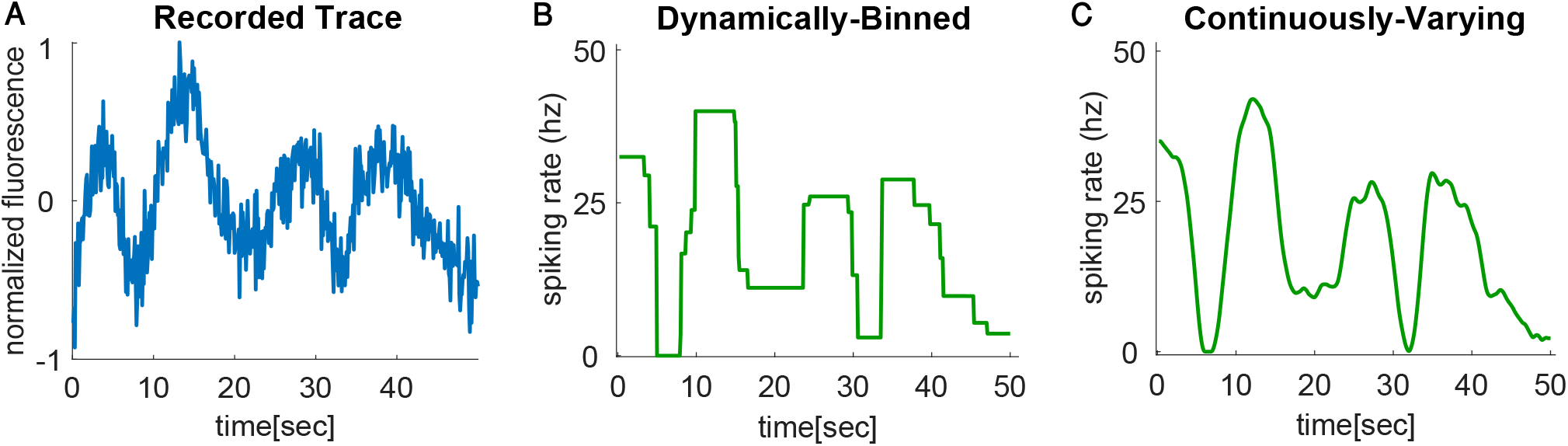
Illustrations of fluorescence trace and spiking rates estimated with two different penalties. A. An example of a recorded fluorescence trace. B. The spiking rate deconvolved from the recorded trace by solving (6) using the *Dynamically-Binned* penalty in (7). C. The spiking rate deconvolved from the recorded trace by solving (6) using the *Continuously-Varying* penalty in (8).

#### Proposition 2.

*The pair* 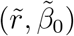 *is a solution to problem* (10) *if and only if* 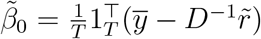 *and* 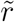 *is a solution to*

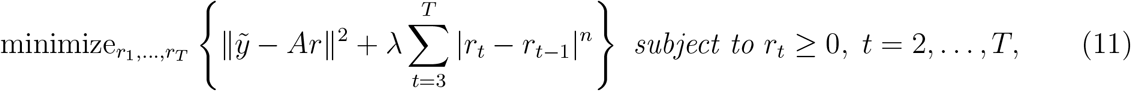

*where* 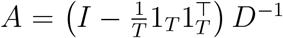 *and* 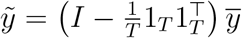, *with I the T × T identity matrix and* 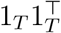 *a T* × *T matrix of ones*.

Note that 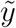 is the mean-centered version of 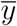, and *A* is the column-mean-centered version of *D*^−1^. We prove Proposition 2 in Appendix A. It implies that we can solve (10) by simply solving (11) to obtain the spiking rate, and then using a closed form expression to obtain the intercept.

With *n* = 1 or *n* = 2, (10) is a convex optimization problem. However, it is not strictly convex and so the solution is not unique. The following proposition indicates that the solution to (10) is invariant under a constant shift in the spiking rate.

#### Proposition 3.

*Let the pair* 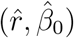 *denote a solution to* (10). *Then the pair* 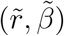 *also solves* (10), *where* 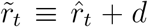 *for t* = 2,…, *T for any* 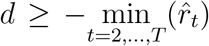 *that satisfies* 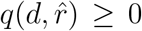, *for* 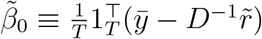 *and a particular choice of* 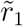.

We provide explicit expressions for 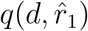 and 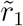 in the proof of Proposition 3 in Appendix B.

## 3 Algorithm

### 3.1 Overview of Proximal Gradient Descent

Equation (11) with *n* ≥ 1 is a convex optimization problem (Boyd & Vandenberghe 2004) that can be efficiently solved for the global optimum. Here, we make use of proximal gradient descent.

We now provide a brief overview of proximal gradient descent; a detailed treatment can be found in Parikh et al. (2014). Suppose that we wish to solve the optimization problem

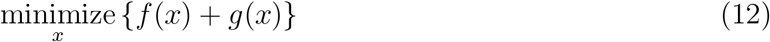

where *f* (·) is a smooth convex function, and *g*(·) is convex but possibly non-differentiable. Then, under mild conditions, an iterative algorithm that initializes *x* at *x*(0), and then at the *t*th iteration applies the update

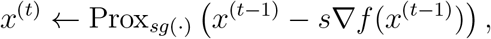

will converge to the global optimum. Here, *s* is a stepsize chosen so that *s* ≤ 1/*L*, where *L* is the Lipschitz constant for the function ∇*f* (·). The notation Prox_*sg*_(·) indicates the *proximal operator* of the function *sg*(·), defined as

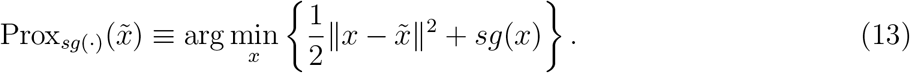

Therefore, proximal gradient descent provides a simple recipe for solving a broad class of convex optimization problems of the form (12), provided that the proximal operator (13) is easily computed, and the function ∇*f*(·) is Lipschitz continuous.

### 3.2 Algorithm for Dynamically-Binned Spiking Rate

We now propose a proximal gradient descent algorithm for solving (10) with *n* =1. By Proposition 2, we can simply solve

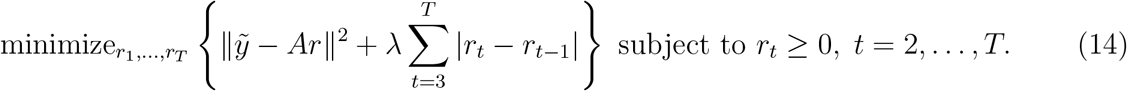

Using the notation of (12), we take 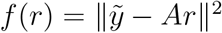 and 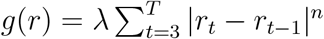.

The following results, which are proven in Appendices C and D, will be useful.

#### Proposition 4.

*The function* 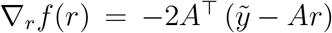 *is Lipschitz continuous with Lipschitz constant L satisfying* 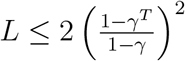.

#### Proposition 5.

*Let 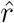 solve the optimization problem*

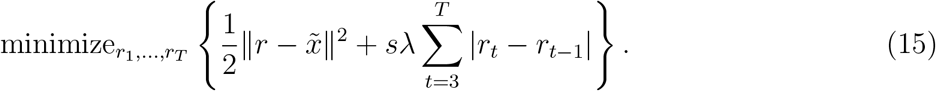

*Then* 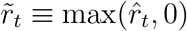 *for t* = 2,…, *T and* 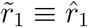 *solves the optimization problem*

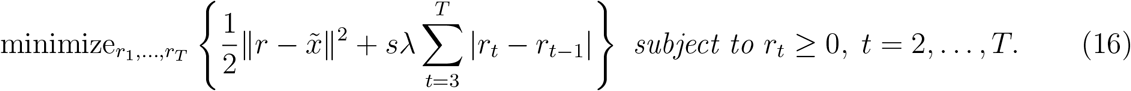

A number of standard solvers, such as the `flsa` solver in `R` (Hoefling 2010), are available to solve (15). We solve (15) by implementing the proposal of Condat (2013). Proposition 5 implies that given a solution to (15), solving (16) is straightforward.

Propositions 2, 4, and 5 lead directly to Algorithm 1 for solving (14).

### 3.3 Algorithm for Continuously-Varying Spiking Rate

We now propose a proximal gradient descent algorithm for solving (10) with *n* = 2. By Proposition 2 it suffices to solve

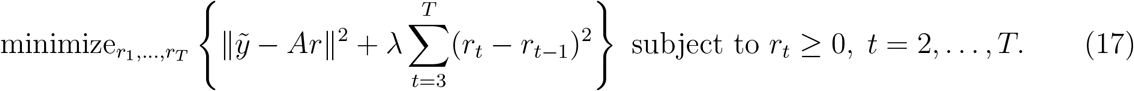

We can further express (17) as

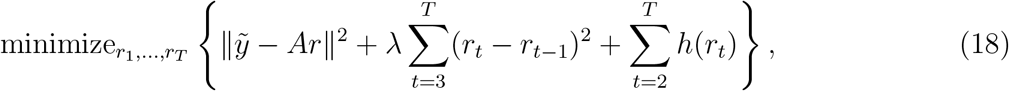

with

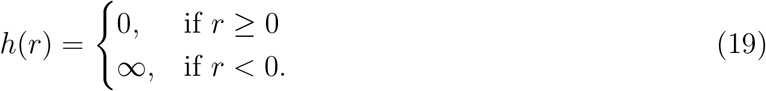

**Algorithm 1.**
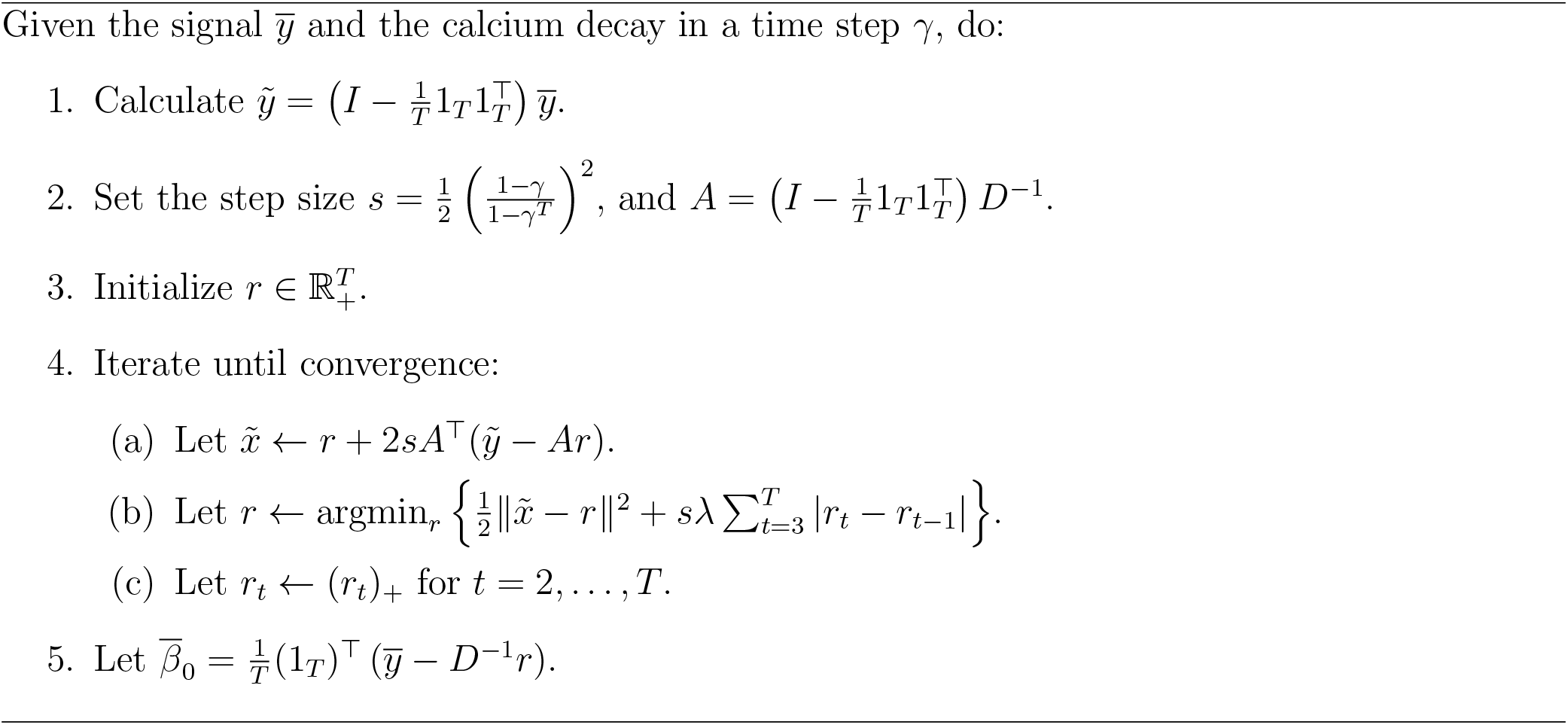
Dynamically-Binned Rate Deconvolution: Solving (10) with *n* =1

This allows us to express the objective function in (17) as *f*(*r*) + *g*(*r*) where

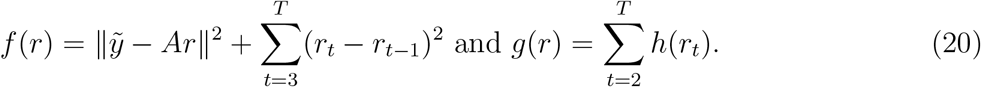

#### Proposition 6.

*The function* 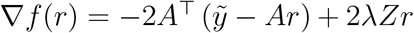 *is Lipschitz continuous with Lips chitz constant L satisfying* 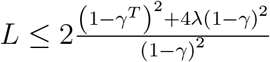, *where*

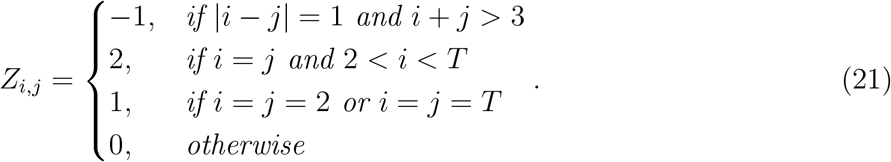

We note that 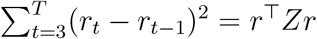. Proposition 6 is proven in Appendix E.

#### Proposition 7.

*For any s* ≥ 0, *the solution to the optimization problem*

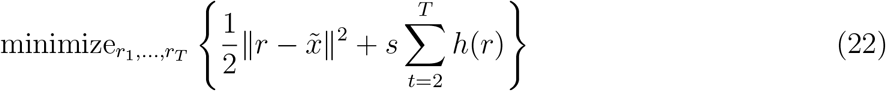

*is given by* 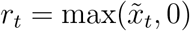 *for t* = 2,…, *T and* 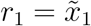.

Proposition 7 is straightforward and the proof is omitted. Propositions 2, 6 and 7 lead to Algorithm 2 for solving (17).

In Appendix F we consider solving (10) with *n* = 2 in the absence of the non-negativity constraint on *r*.

**Algorithm 2.**
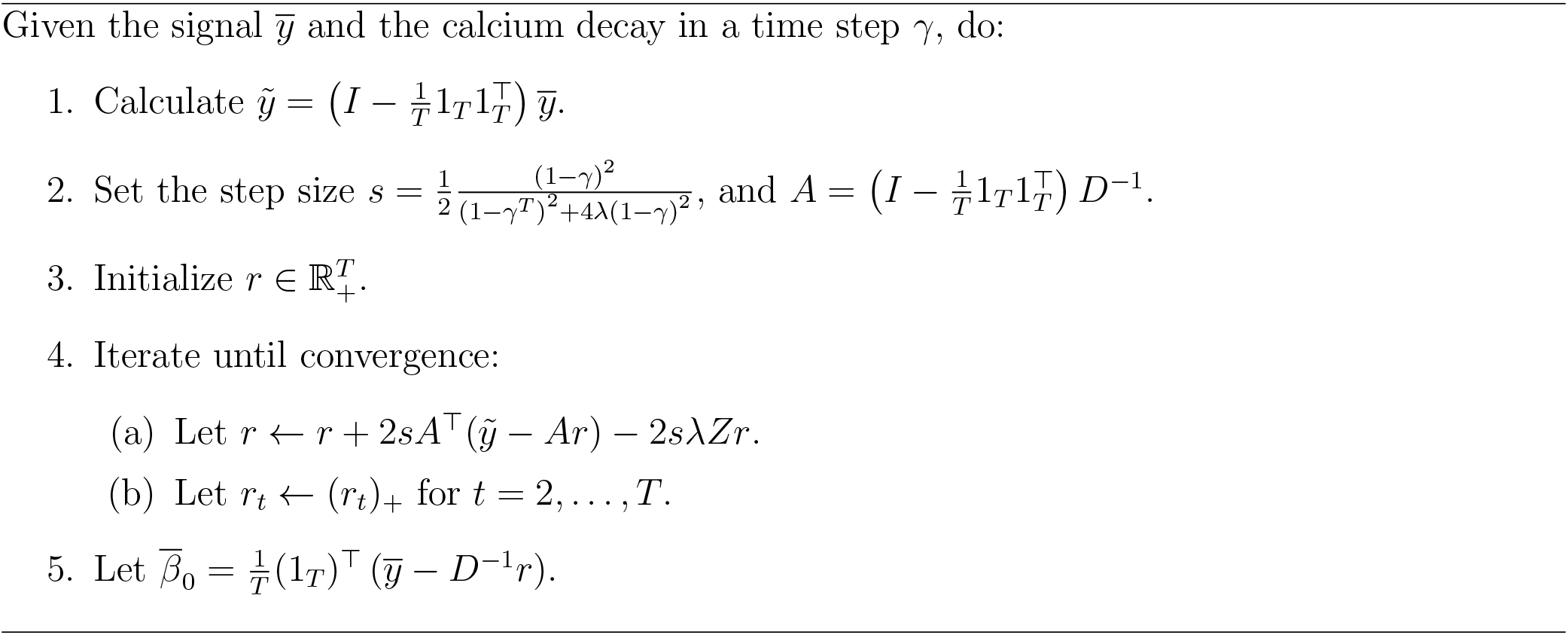
Continuously-Varying Rate Deconvolution: Solving (10) with *n* = 2

## 4 Results on Simulated Data

### 4.1 Evaluation of Algorithms 1 and 2

Here we evaluate the performances of Algorithms 1 and 2 on simulated fluoresence data generated from randomly drawn rate traces. We measure the difference between the true rate *r* and the deconvolved rate *r^est^* as

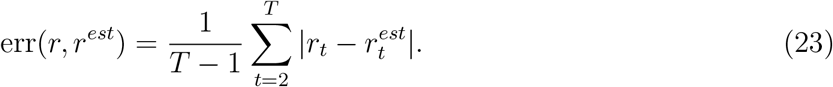

Since the solution to (10) is unique only up to a contant shift (see Proposition 3), we compute (23) after subtracting the mean from both *r_t_* and 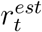 for *t* = 2, …, *T*.

We begin by generating 500 random piece-wise constant rate traces and 500 random continuous rate traces, and use those to calculate calcium and fluorescence traces (details are given in Appendix G; see Figures 2A, 2B, 2D, 2E). We used Algorithms 1 and 2 to deconvolve the fluorescence traces generated from piece-wise constant and continuous rates, respectively.

**Figure 2:**
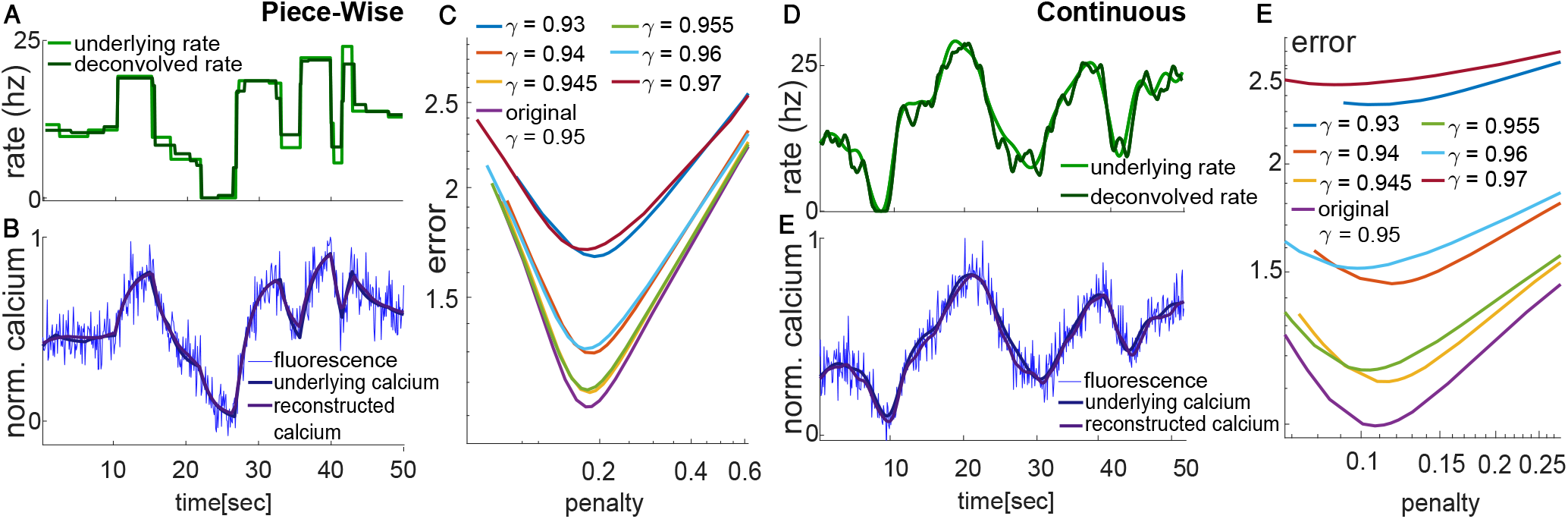
Deconvolution of the fluorescence signal into spiking rate. A–C. Dynamically-binned deconvolution. A,B. Underlying piece-wise constant rate (light green) was created by the random process described in Appendix G. The rate was transformed, using the model (4), into the underlying calcium trace (dark blue). White noise and a constant shift were added to the latter to create the noisy fluorescence signal (blue, plotted here without the shift). Algorithm 1 was used to estimate the spiking rate. The resulting estimated rate (dark green) successfully recovers the underlying rate. The estimated rate was used to estimate the underlying calcium trace (purple) using (4). C. Error of the deconvolution algorithm, using (23) and a range of values of *γ* and λ, averaged over 500 simulated data sets. D-F. Continuously-varying rate deconvolution. Algorithm 2 was used to estimate the spiking rate; details are as in A–C.

For each fluorescence trace, we performed the deconvolution for a range of values of the penalty λ and the decay rate *γ*. For each pair of λ and *γ*, we calculated the average error (23) across all 500 simulated data sets, as well as the average of 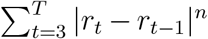. The results are summarized in Figure 2C and 2E.

Our results indicate that deconvolving the rate using the original calcium decay constant γ indeed yields the lowest error for most values of the penalty λ.

### 4.2 Comparisons with Other Methods

We compare our algorithms’ performances to those of other approaches that can be used to analyze wide-field calcium imaging data.

1. *First Differences Deconvolution.* This method assumes that the calcium is represented directly by the fluorescence. Hence, the rate can be found by assuming 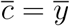 in (4), yielding

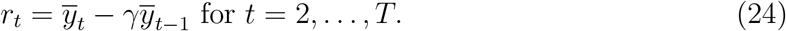 In the special case of *γ* = 1, this method is equivalent to estimating the rate by computing the first differences of the observed fluorescence. We used a moving average window to smooth the resulting estimated rate, and added a mean shift to the result in order to avoid negative rates. This method is a natural and simple alternative to our approach: it bypasses estimation of the underlying calcium, and replaces the penalties in (7) and (8) with a simple *post hoc* smoothing.
2. *Lucy-Richardson Deconvolution*. This iterative algorithm was originally used in astrophysics to restore the light source from a filtered blurred image (Lucy 1974, Richardson 1972). It assumes that the blurring is a result of a convolution by a filter *f* and additive noise. The algorithm maximizes the likelihood of the original image and the denoised image, assuming the noise has a Poisson distribution. In wide-field calcium imaging, this approach has been applied with a one-dimensional filter given by *f_t_* = *γ^t^* (Wekselblatt et al. 2016). To improve performance, we smooth the results using a moving average. The algorithm guarantees non-negativity of the estimated rate.

We compare the above methods and Algorithms 1 and 2 by simulating continuously-varying spiking rates (Appendix G) and measuring the errors (23) of the deconvolved rates. Results are shown in Figure 3 for a range of values of *γ* and λ, averaged over 500 simulated data sets. We also measured the fluctuation in each of the deconvolved rate traces by

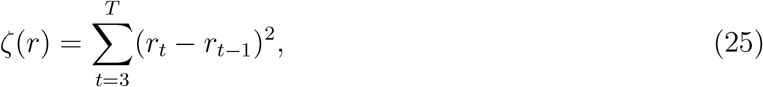

which is the penalty used in (10) with *n* = 2.

**Figure 3:**
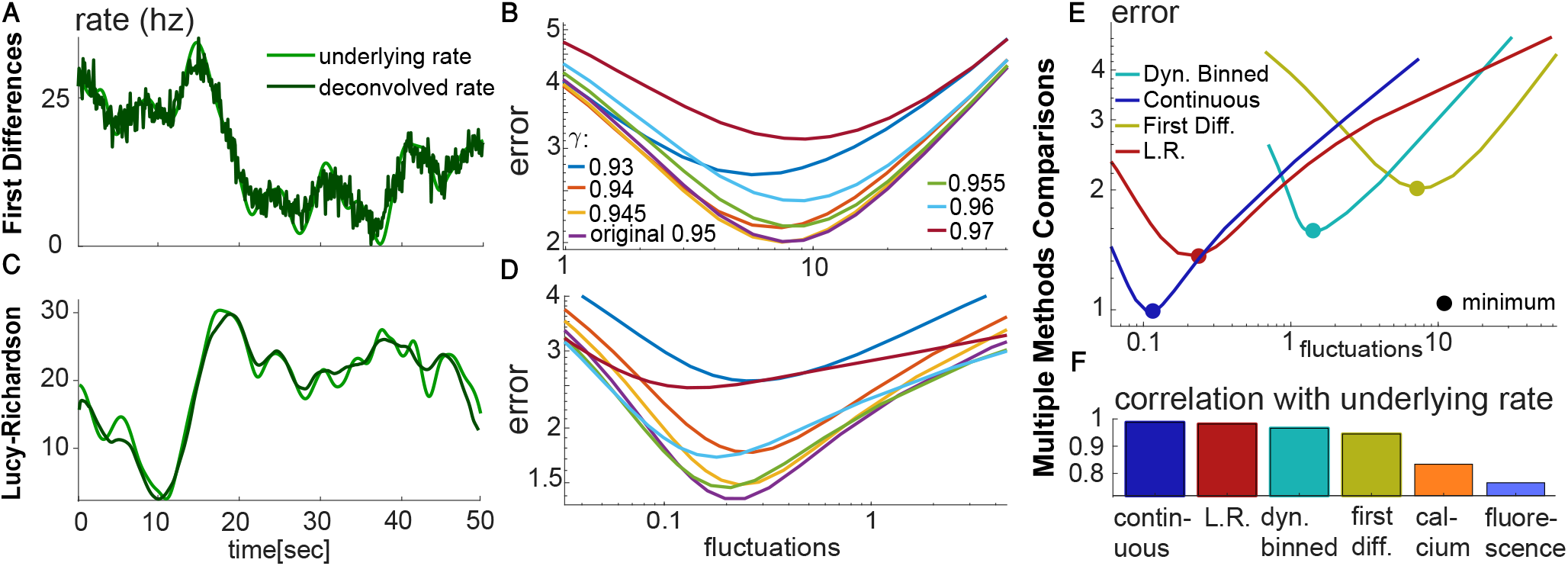
Deconvolution using competing approaches. A-B. Deconvolution of continuously-varying spiking rate using the first differences method. A. An example: Underlying continuous rate (light green) and estimated rate using the first differences method (dark green). The example was generated using the parameter set that yields the lowest average error for this method. B. Error of the first differences method quantified using (23). The x-axis displays the fluctuations in the estimated rate defined in (25). We used a range of *γ* values and a variety of smoothing window lengths. C-D. Similar to A-B but for the Lucy-Richardson (L.R.) method. E. Comparisons of Algorithms 1 and 2 with other methods. F. Correlation of the underlying rate with the deconvolved rate achieved by our algorithms, other methods, or the underlying calcium or fluorescence.

We find that both the first differences and Lucy-Richardson approaches perform best when the true value of *γ* is used (Figures 3B and 3D). However, the best results for the first differences approach are achieved with larger fluctuations (i.e. larger values of the penalty in (25)) compared to other methods (Figures 3B and 3E). This is also evident from inspecting the best deconvolved rate for the example trace (Figure 3A). Lucy-Richardson achieves the best fit with fewer fluctuations (Figure 3D), which results in a smoother deconvolved rate (Figure 3C).

Figure 3E indicates that our continuously-varying spiking rate algorithm performs the best, with the smallest error and the smallest fluctuations at its minimum error (with γ = 0.95 and λ = 2750). Lucy-Richardson comes in second, just ahead of our dynamically-binned algorithm, and far ahead of the first differences method. When we simulated data with a piece-wise constant spiking rate (not shown), our dynamically-binned algorithm performed the best, with Lucy-Richardson second and just ahead of our continuously-varying algorithm, and the first differences method once again far behind.

Last, we calculated the correlations of the deconvolved rate traces, the underlying calcium, and the fluorescence with the underlying rate traces. We see in Figure 3F that all deconvolution methods yield substantially better estimates of the underlying rate than simply using the underlying calcium or observed fluorescence.

## 5 Results on Recorded Data

### 5.1 Whole Dorsal Surface Recordings

Here we evaluate the performances of Algorithms 1 and 2 and the first differences and Lucy-Richardson deconvolution methods on a dataset described in Musall et al. (2018). The dataset consists of fluorescence traces recorded simultaneously at 20Hz from the whole dorsal surface of a genetically encoded GCaMP6s mouse. During the one-hour recording, the mouse performed multiple trials of some task. Dimension reduction (Musall et al. 2018), hemodynamics removal (Wekselblatt et al. 2016), and Δ*F/F* transformation were performed on the traces. Qualitatively similar results are achieved after performing only a Δ*F/F* transformation prior to deconvolution (not shown).

In this data set, because the true spiking rate is unknown, we split each recorded fluorescence trace 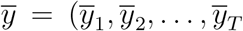 into “odd” and “even” traces, 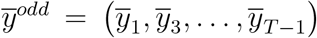 and 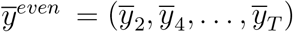. We then deconvolve the odd trace *y^−odd^* to estimate the spiking rate *r^odd^* and the intercept, 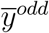. To calculate *D*, defined in Section 2.2, we use *γ* = 0.95, the estimated decay (Chen et al. 2013) for GCaMP6s mice at the 10Hz frequency corresponding to the “odd” and “even” traces.

Next, we apply the second line in (4), adapted to the odd observations, in order to compute 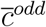 according to 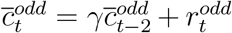 for *t* = 1, 3, 5,…,*T* – 1. We then apply the first line in (4) to compute

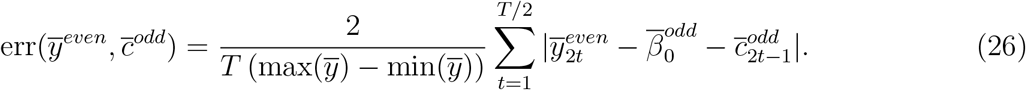

The four different deconvolution methods were applied for a range of tuning parameter values. We find that Algorithm 2 results in the smallest error out of all methods; see Figure 4J. The first differences method as well as Algorithm 1 result in very similar performances, while the latter has less fluctuation in the deconvolved rate (see (25)); this can be seen in Figures 4E-J.

**Figure 4:**
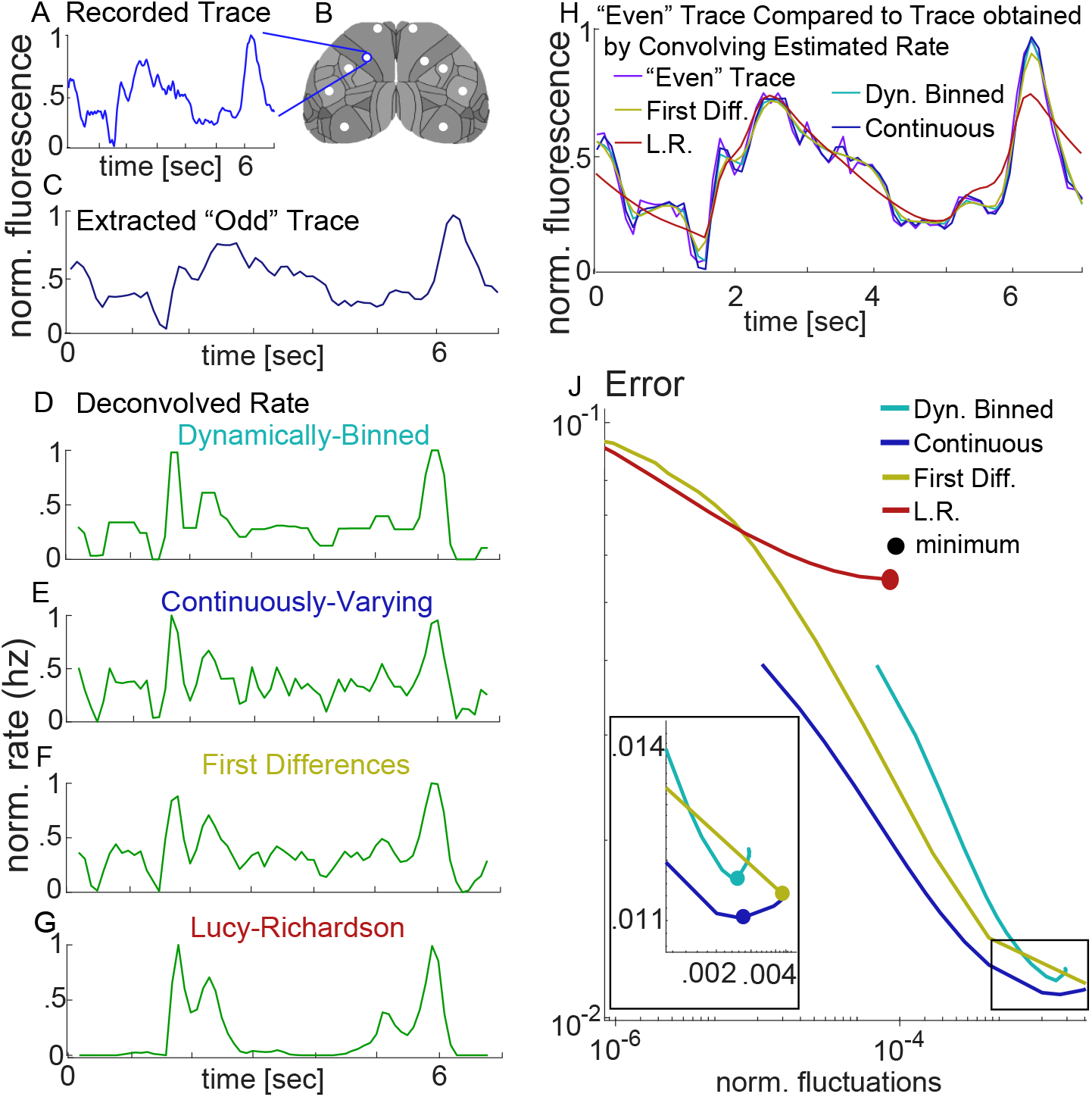
Deconvolution of recorded data from Musall et al. (2018). A. An example of a fluorescence trace recorded from a pixel during a single trial. B. Schematic of a whole dorsal surface wide-field view. White circles indicate the locations of pixels from which fluorescence traces were recorded. The example in A was taken from the pixel circled in blue. C. A fluorescence trace created by extracting from A. the odd time points. D-G. Deconvolution of the fluorescence trace in C. into spiking rate using the D. dynamically-binned algorithm (Algorithm 1), E. continuously-varying algorithm (Algorithm 2), F. first differences method, and G. Lucy-Richardson method. The rates in D-G are displayed with tuning parameters selected to minimize the error in J. H. The “even” trace from A is displayed alongside the results of convolving the spiking rates in D-G obtained from the “odd” trace. J. The relative error between the “even” fluorescence trace and the convolved calcium from the deconvolved spiking rate using the “odd” fluorescence trace (26). The error is displayed as a function of the normalized fluctuations, defined by (25) divided by max(*r^odd^*)^2^. Different fluctuation levels and errors were achieved by varying λ for the dynamically-binned and continuously-varying algorithms, and by varying the smoothing window for the first differences and Lucy-Richardson methods. Larger fluctuation values correspond to lower values of λ and smaller smoothing windows. When λ = 0, no smoothing occurs; this corresponds to the right-hand side of the figure. Results were averaged across traces from all 10 pixels highlighted in B., and 100 trials for each pixel. All fluorescence traces were shifted to have only positive values prior to performing the Lucy-Richardson method.

### 5.2 Parallel Wide-Field and Spike Recordings

Here we compare the performances of Algorithm 1, Algorithm 2, first differences and Lucy-Richardson on parallel recorded spiking data described in Clancy et al. (2019). The data consists of the number of spikes recorded from a multi-channel silicon probe in V1 in each 25ms bin and, simultaneously, a fluorescence trace recorded from the same location at 40Hz. A Δ*F/F* transformation was applied to the fluorescence trace after recording. We note that the probe detects only a subset of the spikes that contribute to the calcium fluorescence. To test the different deconvolution methods, we use the rate of the detected spikes as an estimate of the underlying spiking rate.

To evaluate the methods, we calculated the difference between the deconvolved and the recorded spike rates (23), after dividing each spike rate by its standard deviation.

A decay rate of *γ* = 0.97 has been reported in the literature (Clancy et al. 2019, Chen et al. 2013) for GCaMP6f mice recorded at 40Hz. We investigated a range of values of *γ* and found that all methods attained the lowest error (23) with *γ* = 0.975 except Lucy-Richardson, which attained the lowest error with *γ* = 0.98. We also found that the Lucy-Richardson method attained its lowest error when its fluctuation (defined in (25)) is quite small, Figure 5D. This agrees with results shown in Figures 4G and 4J. The smallest error, across all methods and parameter values, was achieved by the dynamically-binned algorithm, shown in Figure 5D. Comparable error values were achieved by the continuously-varying and Lucy-Richardson methods.

**Figure 5:**
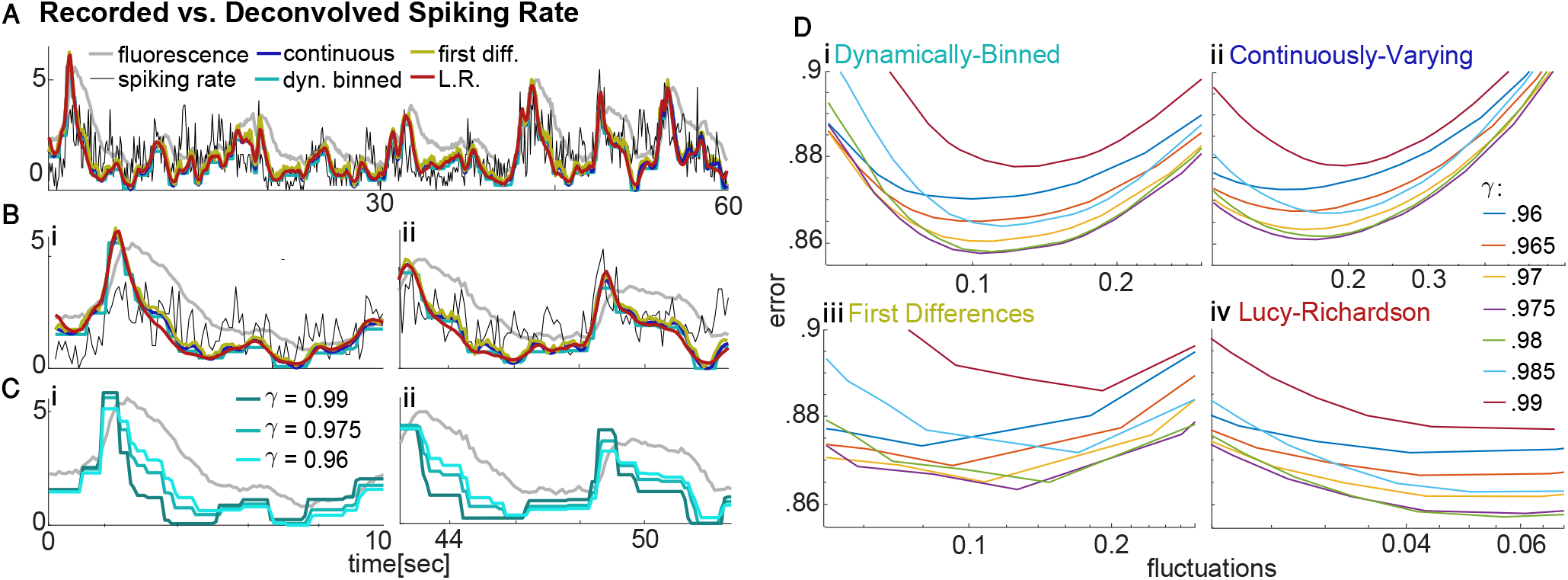
Deconvolved spiking rate compared with recorded spiking rate. A. An example of spiking rate (black) recorded in parallel to the fluorescence (gray), along with the rates estimated by the four deconvolution methods. Tuning parameters were selected to yield the lowest error (23). B. Enlarged regions from A for (i) 0-10 sec and (ii) 42-52 sec. C. Deconvolved spiking rate (shades of turquoise) calculated using different calcium decay values, γ, to deconvolve the fluorescence trace (gray) for (i) 0-10 sec, and (ii) 42-52 sec, used for the dynamically-binned algorithm. D. The error (23) between the recorded spiking rate and the estimated spiking rate obtained using (i) the dynamically-binned algorithm, (ii) the continuously-varying algorithm, (iii) first differences, and (iv) Lucy-Richardson. For each of the methods, the error was calculated for a range of tuning parameter values. Larger fluctuation values, defined in (25), correspond to lower values of λ for Algorithms 1 and 2 and smaller smoothing windows for first differences and Lucy-Richardson methods. Results were averaged across 75 traces, each of which was 60 seconds long.

## 6 Discussion

In this paper, we have proposed two new approaches for estimating the spike rate from wide- field calcium imaging data. The first approach assumes that the true spike rate is piece-wise constant with bins that must be dynamically estimated from the data, whereas the second assumes that the true spike rate varies continuously. We have shown that these approaches outperform existing approaches for spike rate estimation from wide-field calcium imaging data on two data sets. Furthermore, they perform well regardless of whether a simple ΔF/F transformation is performed, or whether hemodynamics removal and dimension reduction are also performed.

In many data sets, multiple fluorescence traces are available from nearby brain regions. The approaches proposed in this paper analyze each fluoresence trace separately, without exploiting the presence of multiple traces. We leave to future work the development of an approach for spike rate estimation that more accurately estimates the spike rate by carefully modeling the spatial dynamics among the fluorescence traces.

## 7 Code Availability

Our code is available at https://github.com/meravstr/Wide-Field-Deconvolution.

## 8 Acknowledgements

We are grateful to Kelly Clancy for providing parallel fluorescence and spiking data recorded in the Thomas Mrsic-Flogel lab. We also thank Simon Musall from Anne Churchland’s lab for providing fluorescence data. MS thanks Nili Guttmann-Beck, Amir Beck Israel Nelken and Matteo Carandini for helpful notes, and OCNC for hosting her during the initiation of this project. DMW has been supported by NSF CAREER DMS-1252624, a Simons Investigator Award in Mathematical Modeling of Living Systems, and NIH grants DP5 0D009145, R01 EB026908, R01 DA047869. ESB was supported by NSF DMS Grants 1514743 and CAREER-1056125. We also acknowledge support from the Sackler Foundation, the Swartz Foundation via the Swartz Center for Theoretical Neuroscience at the University of Washington, and the NSF-ERC Center for Sensorimotor Neural Engineering at the University of Washington. We thank the Allen Institute founders, Paul G. Allen and Jody Allen, for their vision, encouragement and support.

## A Proof of Proposition 2

### Lemma 1.

*Let* 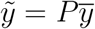 *with* 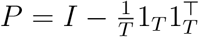 *and* 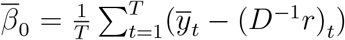. *Then*, 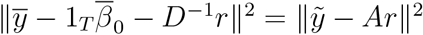.

*Proof of Lemma 1.* Since *P^T^P* = *p* and *P*1_*T*_ = 0, it follows that

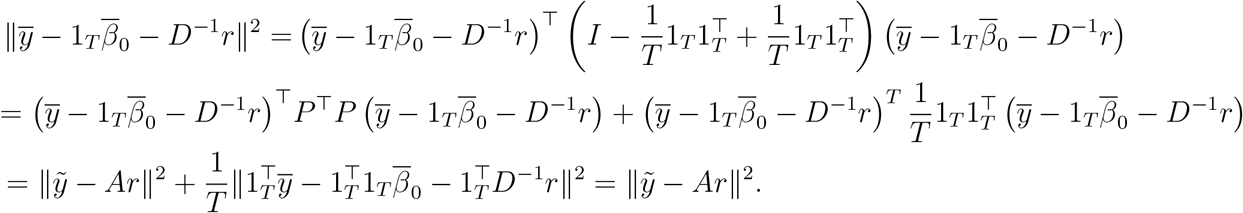

The last equality follows from the fact that 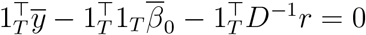.

*Proof of Proposition 2.* Taking the derivative of (10) with respect to 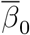 and setting it equal to zero, we find that 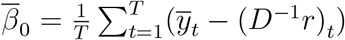. The result follows from Lemma 1.

## B Proof of Proposition 3

*Proof of Proposition 3.* Proposition 2 states that 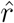 is a solution to (10) if and only if it is also a solution to (11). Hence, f minimizes the objective function 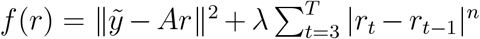. Therefore, any 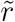 that satisfies 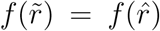 with 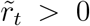 for *t* = 2,…,*T* is a solution to the optimization problem (10) as well.

If we construct 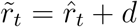 for *t* = 2,…, *T* with 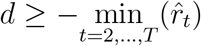, then 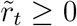 for *t* = 2,…, *T*. In addition, 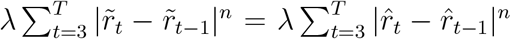. So to show that 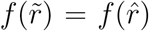, it suffices to show that 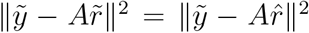. We will do so by choosing an appropriate value for 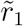. In particular, notice that

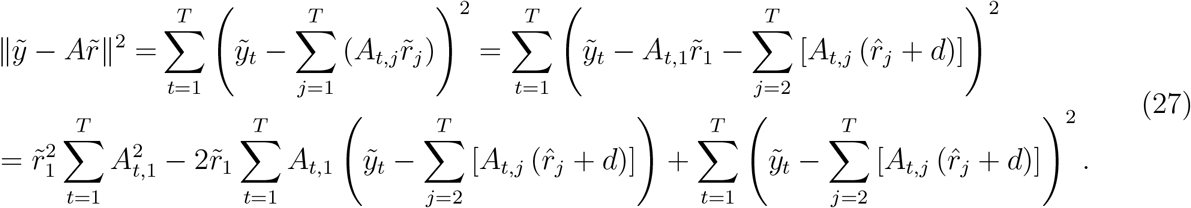

So it suffices to find the value of 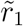 for which (27) equals 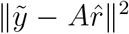. We observe that (27) is quadratic in 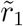. We define

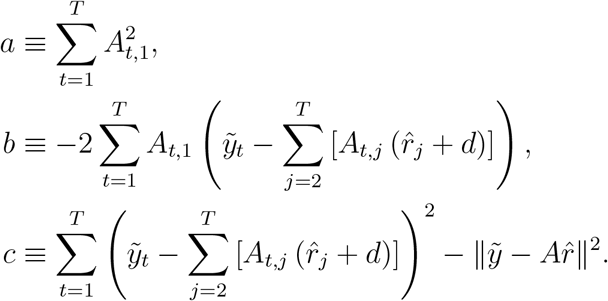

A solution exists provided that *b*^2^ – 4*ac* ≥ 0, in which case 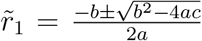. Defining 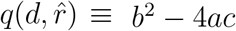, the proof is complete.

## C Proof of Proposition 4

Equation (4) implies that

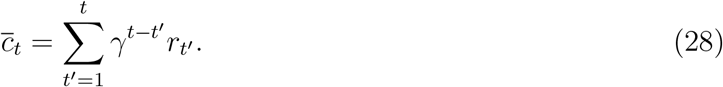

Since 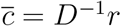, we find from (28) that the entries of *D*^−1^ are given by

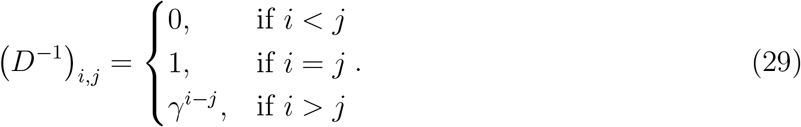

### Lemma 2.

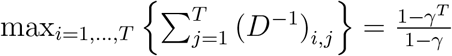.

*Proof.* From (29),

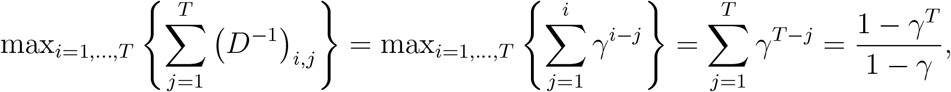

where the last step was calculated using the expression for the sum of a geometric series.

Following the same reasoning, one can also show that 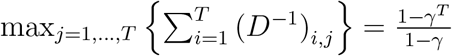.

*Proof of Proposition 4.* The Lipschitz constant of 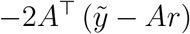 is given by the largest eigenvalue of 2*A*^T^*A* (Bubeck et al. 2015). To find it, it is convenient to first explore the largest eigenvalue of (*D*^−1^)^T^ *D*^−1^, which is a symmetric matrix with positive real entries. By the Perron Frobenius Theorem and Lemma 2, we can bound its largest eigenvalue by its largest single row sum:

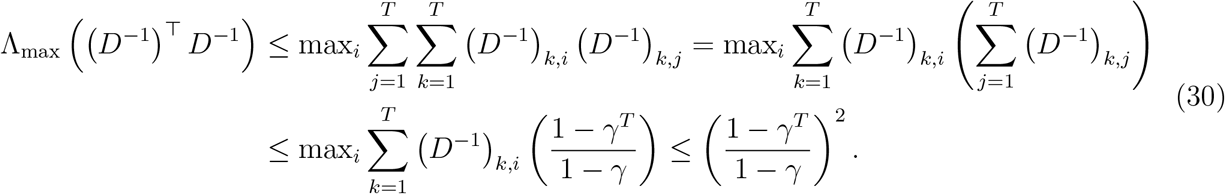

We recall that *A* = *PD*^−1^ and we observe that *P*^T^*P* = *P*, where 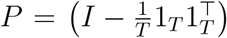, with *I* the identity matrix and 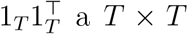 matrix of ones. Together with (30) and Rayleigh quotient properties, the following holds for any vector *u*:

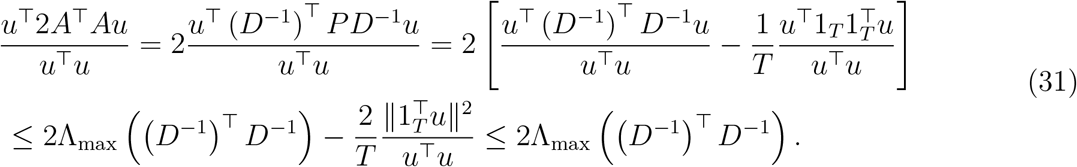

Since the above inequality holds for any vector *u*, including the eigenvector associated with the largest eigenvalue of 2*A*^T^*A*, it follows that

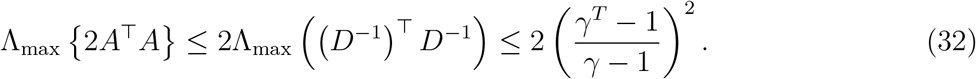

Hence the Lipschitz constant of 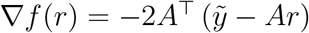 is bounded above by 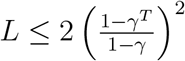.

## D D Proof of Proposition 5

We will make use of the *generalized sign*, the subdifferential of the *ℓ*_1_ norm, defined as

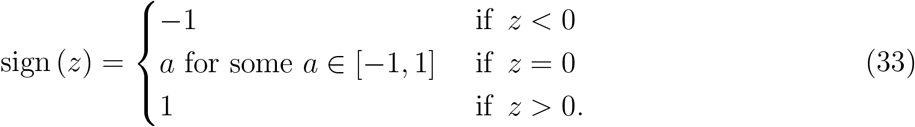

When *z* is a vector, the operation sign(*z*) is applied componentwise.

*Proof of Proposition 5.* The solution 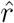 to (15) satisfies the optimality condition

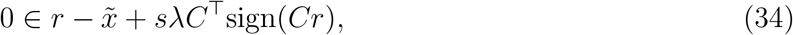

where *C* is a (*T* – 2) × *T* matrix defined as

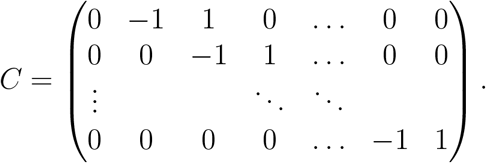

Furthermore, 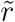 is a solution to (16) if and only if there exists some 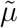 such that the pair 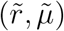 satisfies the Karush-Kuhn-Tucker optimality conditions (Boyd & Vandenberghe 2004), given by

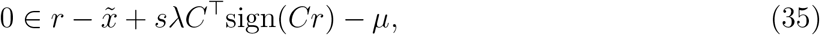

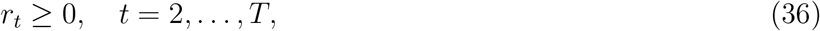

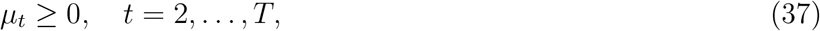

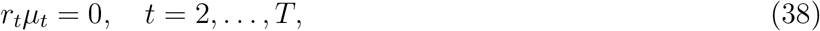

where *μ*_1_ = 0. To complete the proof, we will show that 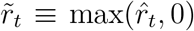 and 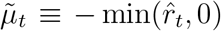 satisfy (35)-(38).

The fact that 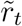 and 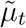 satisfy (36)-(38) follows by inspection. It remains to show that 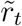 and 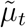 satisfy (35). Notice that 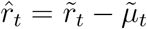. Therefore, because 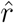 solves (15), it follows directly from (34) that

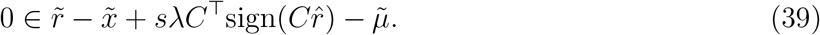

Inspection of the matrix *C* reveals that the elements of 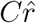 are of the form 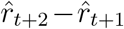. Furthermore, it is straightforward to show that (i) 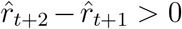 implies that 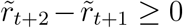; (ii) 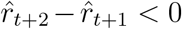 implies that 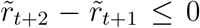; and (iii) 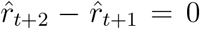 implies that 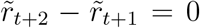. Therefore, 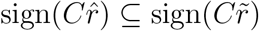. Combining this with (39) directly implies that the pair 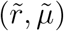 satisfies (35).

## E Proof of Proposition 6

*Proof of Proposition 6.* The Lipschitz constant of the function 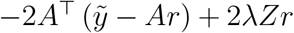 is given by the largest eigenvalue of 2*A*^T^*A* + 2λ*Z* (Bubeck et al. 2015). Since *A*^T^*A* and *Z* are symmetric and real, the largest eigenvalue, Λ_max_, of their sum is bounded by the sum of their largest eigenvalues:

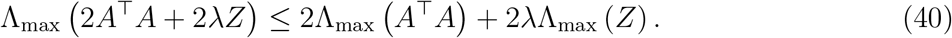

It is not hard to show that Λ_max_ (*Z*) ≤ 4.

Together with (32) and (40), it follows that the Lipschitz constant L satisfies 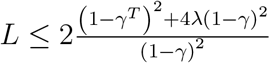.

## F Analytic Solution for the Continuously-Varying Spike-Rate Problem without Non-Negativity Constraints

We consider problem (10) with *n* = 2 and without the non-negativity rate constraint, meaning the problem

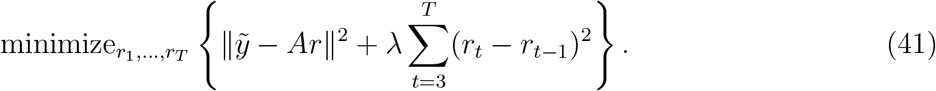

This problem can be solved analytically by computing ∇*f*(*r*) = 0, with 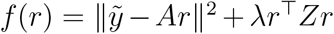 and *Z* defined in (21), yielding:

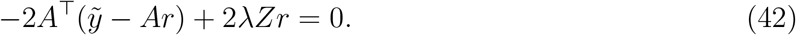

The solution takes the form

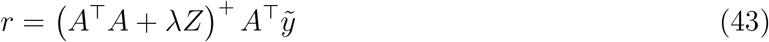

where (*A*^T^*A* + λ*Z*)^+^ denotes the pseudo-inverse of *A*^T^*A* + λ*Z*.

## G Simulation Details

### G.1 Piece-wise Constant Rate

We generated changepoints for the rate from a discrete Unif.(0, 50) distribution. Between each pair of changepoints, we generated the rate from a *N*(0,10^2^) distribution. We then made the rate trace nonnegative by subtracting out its minimum value.

We generated the underlying calcium trace 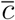 from the rate trace *r* by calculating 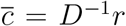, where we used *γ* = 0.95 in constructing *D*^−1^ (29). The value *γ* = 0.95 was estimated as the decay of the calcium in Gcamp6s across 100 milliseconds (Chen et al. 2013).

We made use of only the last *T* = 600 timepoints of the calcium and the rate traces we generated, so that 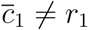, as is the case in a real experiment in which the activity before the first measured point is unknown.

Finally, we generated the observed florescence trace 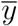 using the model (4) with noise variance 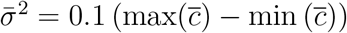, and intercept 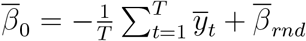, where 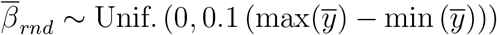.

### G.2 Continuously-Varying Rate

We generated continuous rate traces by integrating the neural network given by the equation 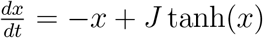, where *x* is a vector of size *N* = 1000, and *J* is an *N* × *N* matrix with i.i.d. entries *J_ij_* ~ *N*(0, 4/*N*). These choices led to chaotic dynamics (Sompolinsky et al. 1988). We sampled the activity of *x*_1_ every 0.5 time steps for 600 steps (resulting in *T* = 1200 time points). To ensure non-negativity of the rate trace, we subtracted out the minimum activity.

Given the rate trace, we then repeated the steps detailed in Appendix G.1 to generate the calcium and fluorescence.

## H Additional Data Analysis

We now re-analyze the fluorescence data of Clancy et al. (2019), originally studied in Section 5.2, without making use of the true spiking rate. This analysis is valuable because in this data set only Δ*F/F* was performed prior to deconvolution. This is in contrast to the fluorescence data of Musall et al. (2018) studied in Section 5.1, on which hemodynamics removal and dimension reduction were also performed.

The results, displayed in Figure 6, generally agree with those found in Figure 4J of Section 5.1. In fact, in a number of other data sets processed using only Δ*F/F*, we found that our algorithms gave good results (results not shown). This suggests that it is not necessary to perform dimension reduction before our algorithms are applied; furthermore, it may be preferable and more natural not to do so.

**Figure 6:**
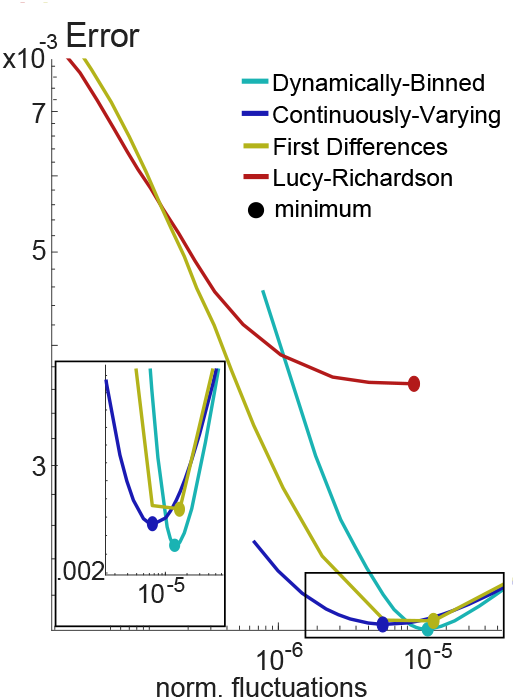
Deconvolution of recorded data from Clancy et al. 2019. This analysis did not make use of the recorded spike rate: only the fluorescence data was used. More details about this figure are provided in Figure 4J.

## References

Aimon, S., Katsuki, T., Grosenick, L., Broxton, M., Deisseroth, K. & Greenspan, R. J. (2015), ‘Activity sources from fast large-scale brain recordings in adult drosophila’, bioRxiv p. 033803.

Allen, W. E., Kauvar, I. V., Chen, M. Z., Richman, E. B., Yang, S. J., Chan, K., Gradinaru, V., Deverman, B. E., Luo, L. & Deisseroth, K. (2017), ‘Global representations of goal-directed behavior in distinct cell types of mouse neocortex’, Neuron 94(4), 891–907.

Boyd, S. & Vandenberghe, L. (2004), Convex Optimization, Cambridge University Press.

Bubeck, S. et al. (2015), ‘Convex optimization: Algorithms and complexity’, Foundations and Trends^®^ in Machine Learning 8(3-4), 231–357.

Chen, T.-W., Li, N., Daie, K. & Svoboda, K. (2017), ‘A map of anticipatory activity in mouse motor cortex’, Neuron 94(4), 866–879.

Chen, T.-W., Wardill, T. J., Sun, Y., Pulver, S. R., Renninger, S. L., Baohan, A., Schreiter, E. R., Kerr, R. A., Orger, M. B., Jayaraman, V. et al. (2013), ‘Ultrasensitive fluorescent proteins for imaging neuronal activity’, Nature 499(7458), 295.

Clancy, K. B., Orsolic, I. & Mrsic-Flogel, T. D. (2019), ‘Locomotion-dependent remapping of distributed cortical networks’, Nature Neuroscience p. 1.

Condat, L. (2013), ‘A direct algorithm for 1-d total variation denoising.’, IEEE Signal Process. Lett. 20(11), 1054–1057.

Friedrich, J., Zhou, P. & Paninski, L. (2017), ‘Fast online deconvolution of calcium imaging data’, PLoS Computational Biology 13(3), e1005423.

Hoefling, H. (2010), ‘A path algorithm for the fused lasso signal approximator’, Journal of Computational and Graphical Statistics 19(4), 984–1006.

Jewell, S., Hocking, T. D., Fearnhead, P. & Witten, D. (2019), ‘Fast nonconvex deconvolution of calcium imaging data’, Biostatistics. URL: https://doi.org/10.1093/biostatistics/kxy083

Jewell, S. & Witten, D. (2018), ‘Exact spike train inference via £_0_ optimization’, The Annals of Applied Statistics 12(4), 2457.

Kalchenko, V., Israeli, D., Kuznetsov, Y. & Harmelin, A. (2014), ‘Transcranial optical vascular imaging (tovi) of cortical hemodynamics in mouse brain’, Scientific Reports 4, 5839.

Lucy, L. B. (1974), ‘An iterative technique for the rectification of observed distributions’, The Astronomical Journal 79, 745.

Makino, H., Ren, C., Liu, H., Kim, A. N., Kondapaneni, N., Liu, X., Kuzum, D. & Komiyama, T. (2017), ‘Transformation of cortex-wide emergent properties during motor learning’, Neuron 94(4), 880–890.

Mann, K., Gallen, C. L. & Clandinin, T. R. (2017), ‘Whole-brain calcium imaging reveals an intrinsic functional network in drosophila’, Current Biology 27(15), 2389–2396.

Masino, S., Kwon, M., Dory, Y. & Frostig, R. (1993), ‘Characterization of functional organization within rat barrel cortex using intrinsic signal optical imaging through a thinned skull’, Proceedings of the National Academy of Sciences 90(21), 9998–10002.

Musall, S., Kaufman, M. T., Gluf, S. & Churchland, A. K. (2018), ‘Movement-related activity dominates cortex during sensory-guided decision making’, BioRxiv p. 308288.

Parikh, N., Boyd, S. et al. (2014), ‘Proximal algorithms’, Foundations and Trends^®^ in Optimization 1(3), 127–239.

Pnevmatikakis, E. A., Soudry, D., Gao, Y., Machado, T. A., Merel, J., Pfau, D., Reardon, T., Mu, Y., Lacefield, C., Yang, W. et al. (2016), ‘Simultaneous denoising, deconvolution, and demixing of calcium imaging data’, Neuron 89(2), 285–299.

Richardson, W. H. (1972), ‘Bayesian-based iterative method of image restoration’, JOSA 62(1), 55–59.

Silasi, G., Xiao, D., Vanni, M. P., Chen, A. C. & Murphy, T. H. (2016), ‘Intact skull chronic windows for mesoscopic wide-field imaging in awake mice’, Journal of Neuroscience Methods 267, 141–149.

Sompolinsky, H., Crisanti, A. & Sommers, H.-J. (1988), ‘Chaos in random neural networks’, Physical Review Letters 61(3), 259.

Swanson, R., Basu, J. & Buzsaki, G. (2018), ‘Investigating hippocampo-cortical dialogue using wide-field calcium imaging and electrophysiology in vivo’, Soc. For Neuroscience Meeting.

Tibshirani, R. (1996), ‘Regression shrinkage and selection via the lasso’, Journal of the Royal Statistical Society: Series B (Methodological) 58(1), 267–288.

Tibshirani, R. J., Taylor, J. et al. (2012), ‘Degrees of freedom in lasso problems’, The Annals of Statistics 40(2), 1198–1232.

Tibshirani, R., Saunders, M., Rosset, S., Zhu, J. & Knight, K. (2005), ‘Sparsity and smoothness via the fused lasso’, Journal of the Royal Statistical Society: Series B (Statistical Methodology) 67(1), 91–108.

Vogelstein, J. T., Watson, B. O., Packer, A. M., Yuste, R., Jedynak, B. & Paninski, L. (2009), ‘Spike inference from calcium imaging using sequential monte carlo methods’, Biophysical Journal 97(2), 636–655.

Wekselblatt, J. B., Flister, E. D., Piscopo, D. M. & Niell, C. M. (2016), ‘Large-scale imaging of cortical dynamics during sensory perception and behavior’, American Journal of Physiology Heart and Circulatory Physiology.

